# Reward-dependent selection of feedback gains impacts rapid motor decisions

**DOI:** 10.1101/2021.07.25.453678

**Authors:** Antoine De Comite, Frédéric Crevecoeur, Philippe Lefèvre

## Abstract

Target reward influences motor planning strategies through modulation of movement vigor. Considering current theories of sensorimotor control suggesting that movement planning consists in selecting a goal-directed control strategy, we sought to investigate the influence of reward on feedback control. Here we explored this question in three human reaching experiments. First, we altered the explicit reward associated with the goal target and found an overall increase in feedback gains for higher target rewards, highlighted by larger velocities, feedback responses to external loads, and background muscle activity. Then, we investigated whether the differences in target rewards across multiple goals impacted rapid motor decisions during movement. We observed idiosyncratic switching strategies dependent on both target rewards and, surprisingly, the feedback gains at perturbation onset: the more vigorous movements were less likely to switch to a new goal following perturbations. To gain further insight into a causal influence of the feedback gains on rapid motor decisions, we demonstrated that biasing the baseline activity and reflex gains by means of a background load evoked a larger proportion of target switches in the direction opposite to the background load associated with lower muscle activity. Together, our results demonstrate an impact of target reward on feedback control and highlight the competition between movement vigor and flexibility.

**Significance statement:** Humans can modulate their movement vigor based on the expected reward. However, a potential influence of reward on feedback control has not been documented. Here we investigated reaching control strategies in different contexts associated with explicit rewards for one or multiple goals, while exposed to external perturbations. We report two strategies: reward could either increase feedback gains, or promote flexible switches between goals. The engagement of peripheral circuits in the modulation of feedback gains was confirmed by the application of a background load that biased feedback vigor directionally, evoking differences in switching behavior in the opposite direction. We conclude that feedback vigor and flexible changes in goal are two competing mechanisms to be selected when interacting with a dynamic environment.

## Introduction

From the toddler picking their favorite toys to the footballer selecting the best path through opponents, humans manifest the exquisite ability to plan and select movements. Movement planning is the process that integrates many task-related factors in order to select the best control strategy for the task (Wong et al., 2015). Amongst these numerous factors, the reward associated with the task induces a modulation of movement vigor in saccadic eye movements (Manohar et al., 2015, 2017) and upper limb reaching movements (Esteves et al., 2016; Summerside et al., 2018). Moreover, recent studies reported that higher reward increases visuomotor responses to disturbances (Carroll et al., 2019) and that the increase of vigor associated with reward is correlated with a reduction of movement variability and an increase in co-contraction (Codol et al., 2020). Together, these previous results suggested an influence of reward on movement planning strategies.

Besides this impact on movement planning, reward also has an influence on movement selection. Indeed, the selection of the best alternative between different options is biased toward movements associated with the highest reward (Trommershäuser et al., 2003, 2008). Similarly, when humans have to select a target, their choice is biased by parameters such as the biomechanical costs incurred when reaching to each potential option, resulting in target selection toward less effortful movements (Cos et al., 2011; Morel et al., 2017).

The commitment to an action actually results from a distributed consensus between low level sensorimotor representations of movement costs (e.g. motor costs) and high level cognitive representations of their outcomes (e.g. reward) (Cisek, 2012). Here we explored the impact of target reward on fast feedback control strategies and tested the distributed consensus theory in a dynamical context by probing the effect of movement reward on feedback control and online motor decisions. Recent studies have sought to investigate whether and how much the factors that characterize action selection during movement planning could also influence movement selection when the hand has already started moving. A first body of work have shown that dynamical changes in target selection can be triggered by mechanical (Nashed et al., 2014) or visual (Kurtzer et al., 2020; Michalski et al., 2020) perturbations occurring during movement. More recently, some studies demonstrated that cognitive factors, such as the reward distribution of a redundant target, also influence online motor decisions (Cos et al., 2021; Marti-Marca et al., 2020). They revealed that the reward distribution of a redundant target influences online motor decisions and suggested a link between the state of the limb (position and speed) at perturbation onset and the outcome of the decision. However, whether the reward of competing alternatives or the level of muscle activity could influence online motor decisions has not been explored yet.

In the present work, we addressed the relationship between target reward and feedback control as well as online motor decisions by applying perturbations while participants performed reaching movements toward one or several targets that differed explicitly by their associated rewards. We hypothesized that the influence of reward on movement planning was linked to the selection of feedback gains, which could impact one’s ability to flexibly change target during movement. In a first experiment, we investigated the influence of reward on feedback control strategies. We then investigated the impact of reward on feedback control when participants had the opportunity to reach to different goals. The goal of the third experiment was to study the competition between feedback gains and the ability to flexibly change movement goal during movement.

We first reproduced previous findings of reward-related increase in velocity toward the target. Importantly, we uncovered that this modulation was associated with an increase in feedback gains and muscle activity. In a second experiment, we observed that the difference in reward between alternative goals could bias online motor decisions and, interestingly, found out that the overall increase in movement vigor was negatively correlated with the potential selection of a new target. Our third experiment confirmed that biases in feedback gains induced experimentally were negatively correlated with the ability to switch goal during movement. These findings demonstrate that movement reward modulates both planning and feedback control, and involves the peripheral motor system through modulation of muscle co-contraction and reflex gains. Moreover, we highlight that this modulation was detrimental to the ability to flexibly switch to a new goal during movement.

## Methods

### Participants

A total of 53 participants were enrolled in this study and took part to one of the three experiments. The first group performed Experiment 1 and included 14 right-handed participants (7 females) ranging in age from 21 to 27. The second group performed Experiment 2 and included 20 right-handed participants (14 females) ranging in age from 20 to 46. The last group performed Experiment 3 and included 19 right-handed participants (11 females) ranging in age from 18 to 52. Participants were naïve to the purpose of the experiments and had no known neurological disorder. The ethics committee of the host institution approved the experimental procedures and participants provided their written informed consent prior to the experiment.

### Experiments

For the three experiments, participants sat on an adjustable chair in front of a Kinarm end-point robotic device (KINARM, Kingston, ON, Canada) and grasped the handle of the right robotic arm with their right hand. The robotic arm allowed movements in the horizontal plane and direct vision of both the hand and the robotic arm was blocked. Participants were seated such that at rest their arm was vertical and their elbow formed an angle of approximately 90 degrees. Their arm was unconstrained and their forehead rested on a soft cushion attached to the frame of the setup. A virtual reality display placed above the handle allowed the participants to interact with virtual targets. A white dot of 0.5 cm radius corresponding to the position of the handle was shown on this display during the whole experiment.

### Experiment 1

In Experiment 1 (Figure 1A top), participants (N=14) were instructed to perform reaching movements to a small circular goal target (1.5 cm radius) located at 25 cm in the y-direction from the start position, a red disk of 1.5 cm radius. Participants had first to put the hand-aligned cursor in the start position, which turned green as they reached it. After a random time delay (drawn from an uniform distribution between 1 and 2s), the goal target appeared as a red disk containing a number (1, 5 or 10) that corresponded to the reward participants would receive if they reached and stabilised within the target for a prescribed time window. Reaction time was not constrained and participants could start the movement whenever they wanted. Following the exit of the start position, participants had up to 600ms to reach the goal target and keep the cursor inside for at least 500ms. The goal target turned green at the end of successful trials, or remained red otherwise. During movements, a mechanical perturbation load could be applied to participants’ hand (33 % of the trials). This load consisted of a lateral step force of ±9 N, with a 10ms linear build-up, aligned with the x-axis. This force was triggered when the hand-aligned cursor crossed a virtual line located at 8 cm from the center of the start position (Figure 1A bottom). Unperturbed and perturbed trials as well as trials with different rewards were randomly interleaved such that participants could not predict the occurrence or the direction of the perturbations. Participants started with a 27-trials training block in order to become familiar with the task and the force intensity of perturbation loads. After completing this training block, they performed 6 blocks of 72 trials interleaved with pauses of three to five minutes to prevent muscle fatigue. Each 72-trials block included: 48 unperturbed trials (16 with each target reward) and 24 trials which contained mechanical perturbations (leftward or rightward, 8 of each reward condition). Participants performed a total of 432 trials, including 24 for each perturbed condition (direction of the mechanical perturbation and value of the target reward). A total score corresponding to the cumulative sum of individual movement rewards was projected next to the goal target. Participants were compensated for their participation according to a conversion of this total score. This conversion was calculated such that each participant received between 10 and 15 € as an incentive to score a maximum number of points during the experiment.

**Figure 1:**
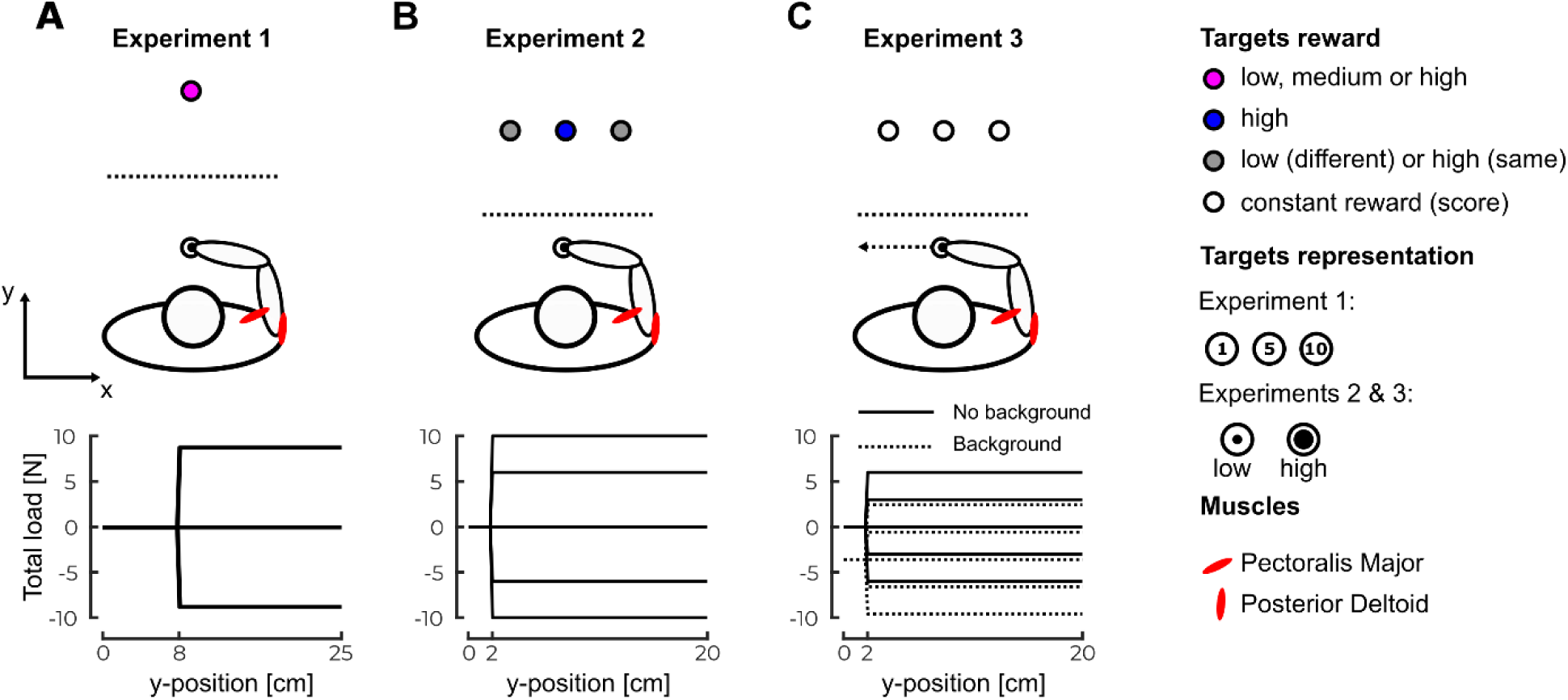
Task paradigms –. **A** Representation of the task paradigm of Experiment 1. Participants controlled a hand-aligned cursor represented by the black dot on a virtual reality display. They had to reach for the goal target, represented by the purple goal target in front of them. This goal target could have a low, medium or high reward (1, 5 or 10 points). The bottom part of the panel represents the load profiles that participants could experience. **B** Representation of the task paradigm of Experiment 2. Participants had to reach for any of the three targets presented in front of them. The central target always had a high reward whereas the two others either had a low or a high reward. The bottom part of the panel represents the perturbation load that participants could encounter during movements. **C** Representation of the task paradigm of Experiment 3. Participants had to reach for any of the three targets presented in front of them. During the second half of the trials a background load force directed leftward was applied prior and during the movement (dashed line bottom panel). The bottom part of the panel represents the possible profiles of the total load forces (perturbation load + background load). EMG data from Pectoralis Major and Posterior Deltoid were collected during all Experiments.

### Experiment 2

Experiment 2 was designed to assess the effect of reward on online motor decisions between competing motor goals. Instead of reaching to a single target, participants (N=20) were instructed to perform reaching movements to any of three circular targets (1.5 cm radius) located at 20 cm in the y-direction from the same start position as in Experiment 1 (Figure 1B top). As in Experiment 1, the goal targets appeared after participants stabilised the hand-aligned cursor in the start position. All three goal targets appeared in each trial, the central one being aligned along the y-axis with the start position and the other two equidistant from this central target at 9 cm along the x-axis. These targets were presented as an inner disk of radius 0.7 or 1.2 cm inside an outer circle of radius 1.5 cm. The purpose of the inner disk was to show the reward associated with the target: the larger the diameter of this disk, the higher the reward. There were two different conditions of reward: either all the targets had the same large reward (*same values* condition) or only the central target had a large reward while the other two had lower rewards (*different values* condition). In a pilot study, we considered a third reward configuration: the central target had a small reward while the other two had higher rewards. We observed that, in this third configuration, the behavior in the absence of perturbation load was biased toward the lateral targets. We therefore decided to exclude this condition to keep the conditions in which participants spontaneously reached for the center target for the largest proportion of trials in the absence of any perturbation load. After a random time delay (drawn from a uniform distribution between 1.5 and 3s), the inner disks of the goal targets turned white and participants had to reach any of these within 400ms to 1000ms to pass the trial. Similar to Experiment 1, the reaction time was not constrained. The trial was successfully completed if participants reached any goal target in the prescribed time window and stabilised the cursor in it for 500ms. The inner disks of the goal targets turned green if the trial was successful and red otherwise. As in Experiment 1, a mechanical perturbation load could be applied to participant’s hand (50 % of the trials, ±6 N or ±10 N, 10 ms build-up aligned with the x-axis Figure 1B bottom). This perturbation was triggered when the hand-aligned cursor crossed a virtual line located at 2 cm from the start position. Unperturbed and perturbed trials as well as trials with different reward distributions and force intensities were randomly interleaved. Participants started with a 58-trials training block followed by 6 blocks of 80 trials. Pauses of three to five minutes were introduced between blocks to prevent muscle fatigue. Each 80-trials block included: 40 unperturbed trials and 40 trials which contained mechanical perturbations. Participants performed a total of 480 trials including 30 trials of each perturbation condition (reward condition and mechanical perturbation condition). Participants were compensated for their participation using the same conversion rule as in Experiment 1.

### Experiment 3

The third experiment was a variant of Experiment 2 and was designed to test the possible impact of muscle activity on online motor decisions by applying a background force orthogonally to the reach path (Figure 1C top). Participants had to perform reaching movements to any of the three targets, located as in Experiment 2. These targets were identical to the large reward target of Experiment 2 and the time course of events in the trial was similar as well except that a leftward background mechanical load of 4N was applied as participants reached the start position and remained on throughout the trials. As in the Experiment 2, a mechanical perturbation load could be applied to participant’s hand during movement (33% of the trials). This load consisted of a ±3 N or ±6 N with a 10ms build-up triggered when the hand-aligned cursor crossed a line located at 2 cm from the start position (Figure 1C bottom) and was added to the background load. Participants first performed a 21-trials training block which did not involve background load. After completing this training, participants performed 4 blocks of 60 trials which did not include the background load. Each 60-trials block included 40 unperturbed trials and 20 trials with mechanical perturbations and they were interleaved with pauses of 3-5min. After these 60-trials blocks, participants performed a second 21-trials training block which included the background load. Once this second training block was completed, participants performed a second set of 4 blocks of 60 trials which included the background load. They thus performed a total of 480 trials amongst which 24 of each condition (with different perturbation loads and background load on or off). To motivate participants, a score corresponding to their number of successful trials was projected next to the goal targets. Participants were compensated a fixed amount for their participation.

### Data collection and analysis

Raw kinematics data was sampled at 1kHz and low-pass filtered using a 4^th^ order double-pass Butterworth filter with cut-off frequency of 20 Hz. Hand velocity along the y-axis was computed from numerical differentiation of the position data using a 4^th^ order centered finite difference.

Surface EMG electrodes (Bagnoli surface EMG sensor, Delsys INC. Natick, MA, USA) were used to record muscles activity during movements. We measured the Pectoralis Major (PM) and the Posterior Deltoid (PD) based on previous studies (Crevecoeur et al., 2019; De Comite et al., 2021) that showed that in this configuration they are stretched by the application of forces opposite to their action, and therefore largely recruited by the feedback responses. Before applying the electrodes, the skin of the participant was cleaned and abraded with cotton wool and alcohol. Conduction gel was applied on the electrodes to improve the quality of the signals. The EMG data were sampled at a frequency of 1kHz and amplified by a factor of 1000. A reference electrode was attached to the right ankle of the participant. Raw EMG data from the pectoralis major (PM) and posterior deltoid (PD) were band-pass filtered using a 4^th^ order double-pass Butterworth filter (cut-offs: 20 and 250 Hz), rectified, aligned to force onset and averaged across trials or time windows as specified in the Results section. The time windows selected for the temporal averaging are the short-latency (20-50ms), the long-latency (50-100ms) and the voluntary time epochs (100-180ms) as proposed in previous work (Pruszynski et al., 2008; Pruszynski & Scott, 2010)

EMG data were normalized for each participant to the average activity collected when participants maintained postural control at the start position against a constant force of 9 N. Data from the pectoralis major were normalized by the EMG activity in the same muscle while performing postural control against a rightward force whereas data from the posterior deltoid were normalized by the EMG activity in the same muscle while performing postural control against a leftward force. This calibration procedure was applied after the second and the fourth blocks in the first two experiments and after the first, third, fifth and seventh blocks in the third experiment. Data processing and parameters extractions were performed using Matlab 2019a.

In Experiment 1, we fitted linear mixed models to determine the effect of the target reward on the kinematics and EMG activity. These models were fitted using the *fitlme* function of Matlab and the formula used was the following:

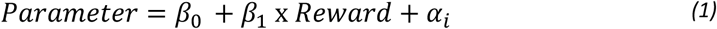

The fixed predictors were the intercept (*β*_0_)and the reward condition (*β*_1_)while the participants were included as a random offset (*α*_*i*_). For all linear mixed model analyses that we performed, we reported the estimate for *β*_1_, the t-statistics for this estimate as well as the corresponding p-value and the *r*^2^of the model. One-tailed paired t-tests were used for post-hoc analyses where we collapsed data across trials and participants to compare the different conditions. Effect size for these tests were reported using Cohen’s d defined as the difference between the means of the two populations divided by the standard deviation of the whole sample.

To analyse the data from Experiments 2 and 3, we designed a multilinear logistic regression model to infer the effect of reward distribution and background load on target choice as the dependent variable, respectively. Considering that the dependent variable was a discrete variable (the chosen target), we use the following logistic regression model:

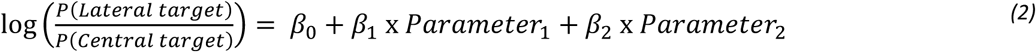

where the first effect (*β*_1_) was the reward condition (Experiment 2) or the presence of a background load (Experiment 3) and the second effect (*β*_2_)was the intensity of the perturbation load. For these logistic regressions, we reported the estimates for *β*_1_ and *β*_2_, their corresponding t-statistics as well as their p-value. For post-hoc analyses in Experiment 2, we used a one-tailed Wilcoxon signed ranked test for which we reported the ranksum, the z-statistics when provided, the p-value as well as the effect size given by the Cohen’s d as defined above. In order to investigate the asymmetry in the parameters *β*_1_ obtained in Experiment 3, we used bootstrap resampling on the individual data to generate 1000 estimates of the *β*_1_ parameter for each condition (leftward perturbation versus rightward perturbation) using the multilinear logistic regression described above. We then assessed the asymmetry of the effect by investigating whether the 95% confidence interval of the difference between these two *β*_1_ parameters contains 0 (Efron, 1979).

In order to determine the effect of the background load on the baseline muscle activity in Experiment 3, we fitted a linear mixed model with interaction terms following this equation:

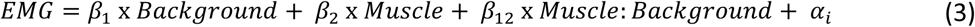

Where the first term (*β*_1_)refers to the background condition, the second (*β*_2_) to the muscle, the third one (*β*_12_) to the interaction term and the last one (*α*_*i*_) to the random offset of participants. For all these *β*, we reported their estimated value as well as their t-statistics, associated p-value and the *r*^2^ of the model. Significance was considered at the level of p=0.05 even though we decide to exactly report any p-value that was larger than p=0.005 as previously proposed (Benjamin & Berger, 2018). In the figures, we reported significant differences for the level p<0.05 (*), p<0.01(**) and p<0.005(***).

## Results

### Influence of the target reward on feedback corrections during movement

To determine whether target reward influences feedback corrections during movement, participants were instructed to perform reaching movements to a goal target associated with a reward that could change across trials (see Methods). During movements, mechanical perturbation loads could be applied to reveal feedback corrections. The occurrence of feedback corrections was assessed by looking at movement kinematics and EMG responses of the muscles stretched by the perturbations.

The mean hand path trajectories across participants are represented in Figure 2A for the different perturbations and reward conditions. Consistent with previous work (Shadmehr et al., 2016; Summerside et al., 2018) we observed a significant increase in forward peak velocity with increasing reward values. Figure 2B shows the differences in the forward velocities between the high (dash-dot lines) or medium (full lines) and low reward conditions unperturbed (top) and perturbed (bottom) trials. The peak forward velocity (defined as the velocity component aligned with the main reaching direction) increased with increasing reward value both for unperturbed (Figure 2D top, linear mixed model: *β*_1_=0.013, t=6.51, p<0.005, *r*^2^=0.76) and perturbed (Figure 2D bottom, linear mixed model, right: *β*_1_=0.014, t=3.76, p<0.005, *r*^2^=0.78, left: *β*_1_=0.018, t=4.68, p<0.005, *r*^2^=0.79) trials. Post-hoc comparisons between low and high reward conditions revealed a significant increase of peak velocity with reward for all perturbation conditions (one-tailed paired t-tests, unperturbed: t=-7.48, p<0.005, d=0.12, left: t=-5.37, p<0.005, d=0.16 and right: t=-3.99, p<0.005, d=0.13). We did not observe any modulation of the reaction time required to initiate movement with the reward (linear mixed model p>0.05) since we instructed participants to initiate movements whenever they wanted (see Methods).

**Figure 2:**
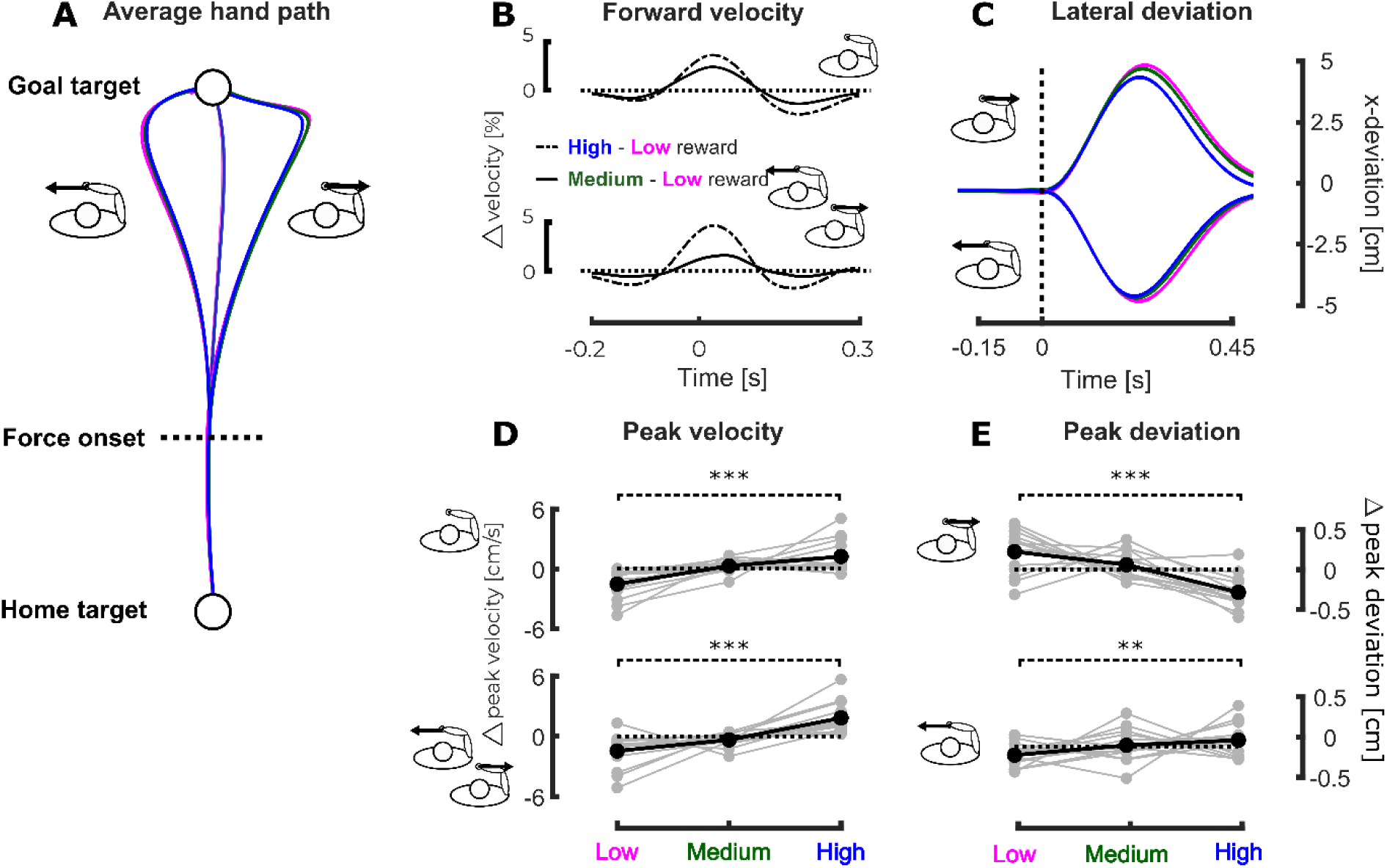
Experiment 1, kinematics – **A** Mean hand path across participants for the different conditions of the first experiment. The purple, green and blue traces respectively correspond to the low, medium and high reward conditions. The dashed line represents the onset of the mechanical perturbation. **B** Mean difference in forward velocity between the high and low (dash-dot line) and medium and low (full line) for the unperturbed (top) and perturbed (bottom) trials. The time axis is aligned on the force onset. **C** Mean hand deviation across participants for the perturbed trials. The hand deviation has been obtained by subtracting the mean hand path to the perturbed hand path in the same reward condition for every subjects. The top part of the graph represents the trials perturbed to the right whereas the bottom part of the graph represents the trials perturbed to the left. **D** Group mean (black) and individual means (grey) of the differential forward peak velocity for the unperturbed trials (top) and perturbed trials (bottom) as a function of the reward condition with respect to average forward peak velocity. **E** Group mean (black) and individual means (grey) of the difference in hand deviation with respect to the mean hand deviation for leftward (top) and rightward (bottom) perturbation in the three reward conditions with respect to the average hand deviation. p<0.05 (*), p<0.01(**), p<0.005(***)

The effect of the mechanical perturbation on the movement kinematics was also dependent on the reward value. Indeed, the lateral hand deviation induced by the mechanical perturbation (Figure 2C), computed as the difference between the hand paths in the perturbed conditions and the mean hand path in the corresponding unperturbed reward condition for each participant, depended significantly on the reward condition. For both perturbation directions (Figure 2E top for leftward and bottom for rightward perturbations), we observed a significant decrease in the maximal hand deviation along the x-axis with increasing reward value (linear mixed models, right: *β*_1_=-0.0025, t=-4.98, p<0.005, *r*^2^=0.38 and left : *β*_1_=-0.0009, t=-2.25, p=0.024, *r*^2^=0.35). Post-hoc comparisons between low and high reward conditions revealed a significant decrease for both perturbation directions (one-tailed paired t-tests, right: t=5.31, p<0.005, d=0.14 and left: t=2.34, p=0.009, d=0.3).

Based on these kinematics analyses and previous studies showing that faster movements and smaller hand deviations induced by perturbations are correlated with high EMG activity (Crevecoeur et al., 2019) we hypothesised that the EMG activity in PM and PD during movement scaled with increasing reward. We investigated this effect both for baseline activity measured during unperturbed trials and for feedback responses to perturbation loads.

We observed a positive correlation between the EMG activity during unperturbed trials and the value of the target reward. Figure 3A represents the mean EMG activity collapsed across muscles and participants for unperturbed trials while the differences between these collapsed EMG activities in the high (dash-dot line) or medium (full line) and the low reward condition are represented in Figure 3B. We binned the EMG activity of each trial in a time bin ranging from 0 to 200 ms after perturbation onset (gray rectangle in Figure 3A and 3B) and fitted a linear mixed model (see Methods) on these binned values to determine whether reward had an influence on the EMG activity (deviations from the mean binned EMG activity in the different reward conditions are represented in Figure 3C and 3D for PM and PD respectively). We observed an increase in EMG activity with the reward in both muscles (PM: *β*_1_=0.028, t=4.603, p<0.005, *r*^2^ = 0.68, PD: *β*_1_= 0.053, t=5.98, p<0.005, *r*^2^=0.66). Post-hoc analyses performed on individual data showed that EMG activity was larger in the high reward condition than in the small one for both muscles (one-tailed paired t-tests: pectoralis, t=4.14, p<0.005, d=-0.118 and deltoid, t=-2.92, p=0.0059, d=0.1653).

**Figure 3:**
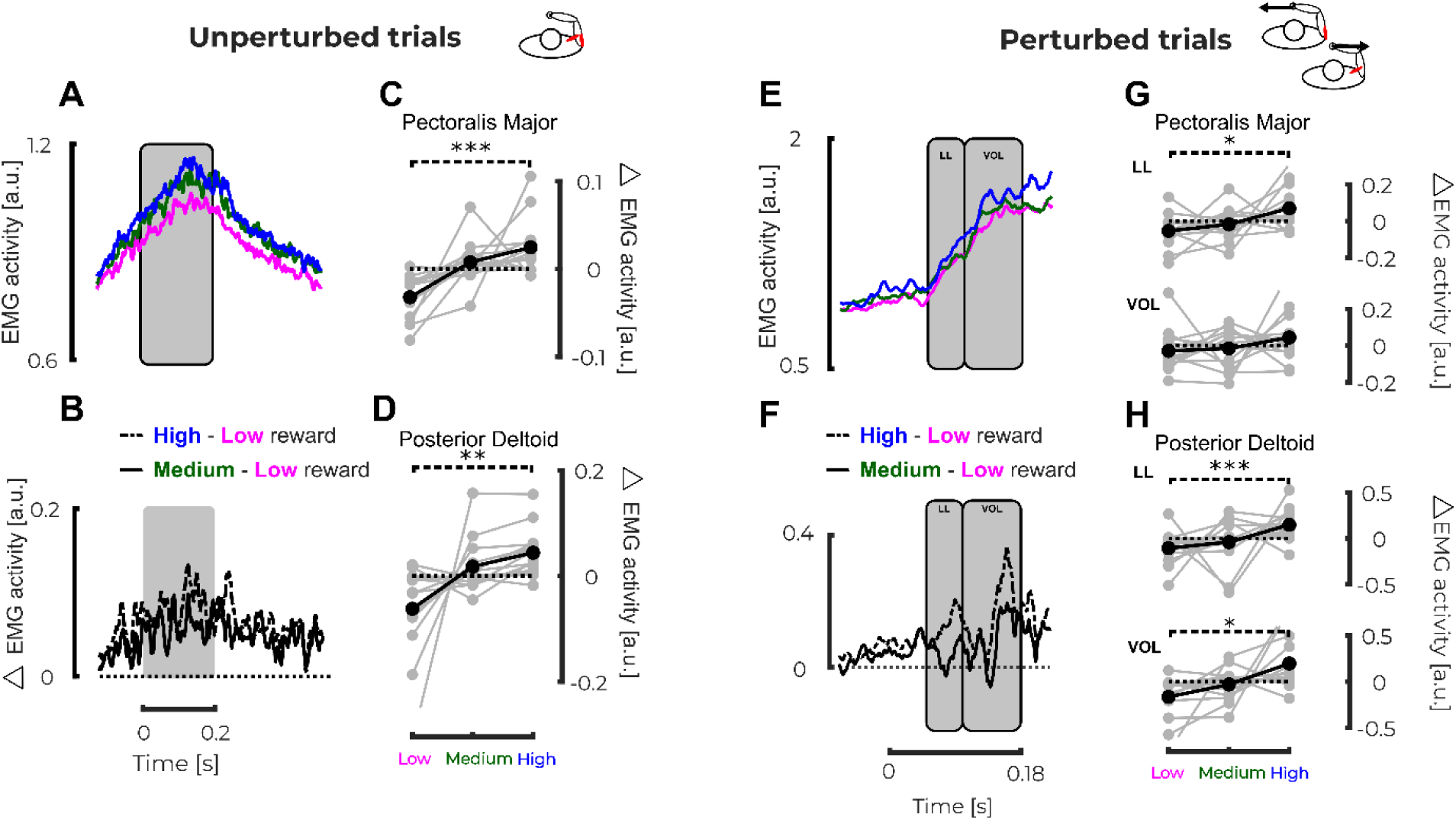
Experiment 1, EMG activity – **A** Mean EMG activity collapsed across muscles and participants for unperturbed trials. The time axis is aligned on force onset. **B** Mean differences in EMG activity collapsed across muscles and participants between high and low (dash-dot line) and medium and low (full line) reward conditions for unperturbed trials. **C** Group mean (black) and individual means (grey) Pectoralis Major EMG activity binned between 0 and 200 ms after force onset for unperturbed trials. **D** Group mean (black) and individual means (grey) Posterior Deltoid EMG activity binned between 0 and 200 ms after force onset for unperturbed trials. **E** Mean EMG activity collapsed across muscles and participants while responding to perturbations. Only the activities of the agonist muscles (ie. PM for rightward perturbation and PD for leftward perturbation) were used. **F** Mean differences in EMG activity collapsed across muscles and participants between high and low (dash-dot line) and medium and low (full line) reward conditions for agonist muscles in presence of perturbation load. **G** Group mean (black) and individual means (grey) differential EMG activity in PM binned in the long latency (50-100 ms, top) and voluntary epochs (100-180 ms, bottom) as a function of the reward condition. **H** Group mean (black) and individual means (grey) of the differential EMG activity in PD binned in the long latency (50-100 ms, top) and voluntary epochs (100-180 ms, bottom) as a function of the reward condition. p<0.05 (*), p<0.01(**), p<0.005(***)

The EMG response to mechanical perturbation in the agonist muscles was also modulated by the reward value. Indeed, linear mixed model analyses performed on the responses measured in PM and PD, when respectively a rightward or leftward perturbation occurred, showed a significant increase of EMG activity with increasing reward in the long-latency epochs (50-100 ms). We reported the EMG activities collapsed across muscles and participants in Figure 3E as well as the difference in these activities between the high (dash-dot line) or medium (full line) and the low reward condition in Figure 3F. For each perturbation direction, we binned the EMG activity of the stretched muscle in the long latency (LL 50-100 ms after force onset) and voluntary (VOL 100-180 ms after force onset) epochs. Figure 3G and 3H respectively represent the deviation from the mean binned EMG activity in these two time bins (LL top and VOL bottom) for PM and PD in the different reward conditions. In PM we observed a significant increase in the LL window (mixed model: *β*_1_=0.0615, t=2.89, p<0.005, *r*^2^=0.64) but even though a positive tendency emerged in the VOL window, no significant increase was observed (mixed model: *β*_1_=0.036, t=1.616, p=0.106, *r*^2^=0.69). Individual pairwise post-hoc comparisons between low and high conditions confirmed these findings (one-tailed paired t-tests: LL, t=-2.48, p=0.0137, d=0.1592 and VOL, t=-1.18, p=0.128, d=0.08). The same holds for PD in which we found a significant increase of EMG activity in LL window with the reward (mixed model: *β*_1_=0.216, t=2.12, p=0.034, *r*^2^=0.63) but no significant effect in the VOL window (mixed model: *β*_1_=0.181, t=1.887, p=0.059, *r*^2^=0.77). In this case however, the individual pairwise comparisons between low and high conditions revealed a significant increase in both time windows (one-tailed paired t-tests: LL, t=-3.68, p<0.005, d=0.11 and VOL, t=-2.57, p<0.01, d=0.07).

An interesting question is whether the effect of target reward reported here could only be attributable to higher movement speed. In other words, could it be that the impact of a higher reward is an increase in movement speed that would therefore modulate the behaviour. To answer that question, we compared the lateral hand deviation observed in trials that have similar peak velocity and investigated whether, in these trials, the reward condition modulates the lateral hand deviation. We performed a linear mixed model analysis on the absolute values of these lateral hand deviations and reported an effect of the reward condition: smaller deviations for higher reward value (*β*_1_=-0.0011, t=-2.61 and p=0.009). This result confirms that reward does not only modulate movement vigor but also the feedback responses to mechanical perturbations.

Therefore, the results of Experiment 1 revealed that the value of the target reward influenced both the movement kinematics and the EMG activity recorded during movement. Indeed, we showed that the hand deviation induced by mechanical perturbations decreased with increasing reward value for both rightward and leftward perturbations. Moreover, the forward peak velocity of reaching movement increased with increasing reward value. Finally, EMG activity in both PM and PD increased with increasing reward value for unperturbed trials and in the long-latency response window for perturbed movement when the muscles were stretched by the perturbation. The modulation of forward hand velocity and baseline EMG activity which also produced increases in feedback responses to perturbation loads was consistent with an increase in control gains previously observed in uncertain dynamical contexts, which was interpreted as a robust control strategy (Crevecoeur et al., 2019).

### Influence of the reward of the different options on online motor decisions

In Experiment 1, we showed that reward of the goal target had an influence on the way humans perform reaching movements to this target similarly to other task parameters such as target shape, presence of obstacles, etc. Moreover, previous studies have shown that these task parameters that modify the control strategies could also influence online motor decisions (Nashed et al., 2014). We therefore designed a second experiment to determine whether reward could also influence online motor decisions. In this second experiment, participants had to reach to any of three potential targets aligned orthogonally to the main reaching direction (see Methods). The central target always had a large reward while the two lateral targets could either have lower reward or a reward equal to that of the central target. We assessed the effect of the difference between central and lateral rewards on online motor decisions by investigating the frequencies of reaching for the lateral targets. Mechanical perturbations that could occur during movement were used to evoke changes in goal target. Because perturbations were unpredictable, a change in reaching frequency for the lateral targets dependent on the perturbation load was indicative of a perturbation-mediated change in goal that occurred during movement. The biomechanical and EMG states at perturbation onset in the same condition were also investigated to determine whether they had an influence on the future decision.

First, we observed that the reward of the lateral targets had a clear effect on the frequency of trials that ended on these targets. Figure 4A represents the hand paths of a typical participant toward the different targets in various conditions. In general, in the absence of perturbation, subjects tend to reach to the central target except for some trials (<1% in the *different values* condition and 8 % in the *same values* condition). In all cases, the frequency of lateral target increased with the magnitude of the perturbation (top and bottom part of Figure 4A for *same* and *different values* conditions respectively). In addition, there was a significant effect of the lateral targets reward on the frequency of lateral target reach: lower frequencies for different rewards. In order to determine the significance of these effects, we identified for each trial the target that was reached at the end of the movement and fitted a multilinear logistic regression on these data to determine whether the reward condition and the force had an influence on the target reached (see Methods). We observed a significant effect of the reward for both the left (*β*_1_ = 1.103, *t* = 10.84, *p* < 0.005) and right (*β*_1_ = 1.666, *t* = 18.45, *p* < 0.005) targets versus the central one. These positive values indicate that the reach proportion to the lateral targets is larger in the same than in the different condition (see Figure 4B and 4C for the *same values* and *different values* conditions respectively). The intensity of the perturbation loads also had a significant effect for both lateral targets versus the central one (left: *β*_1_=-1.33, t=-26.82, p<0.005 and right: *β*_1_=1.25, t=29.07; p<0.005). Due to the sign of the force in the regression model, in both cases the frequency of lateral target reach increased with the force magnitude in absolute value. Post-hoc analyses performed at fixed force levels showed a significant effect of the reward condition on the reaching proportion for all the perturbed conditions. We observed a smaller reach proportion to the left target in the *different values* condition compared to the *same values* condition for both perturbation directions (one tailed Wilcoxon signed-rank test: ranksum=3, p<0.005, d=0.61 - see Figure 4D - and ranksum=1, p<0.005, d=0.49 – see Figure 4E - for loads of -10 and -6N respectively). The mirror effect was observed for the right target: a decrease in the reach proportion in the *different values* condition for both perturbation directions (one-tailed Wilcoxon signed-rank test: ranksum=0, p<0.005, d=0.74 – see Figure 4F - and ranksum=3, p<0.005, d=0.78 – see Figure 4G - for loads of 6 and 10 N respectively).

**Figure 4:**
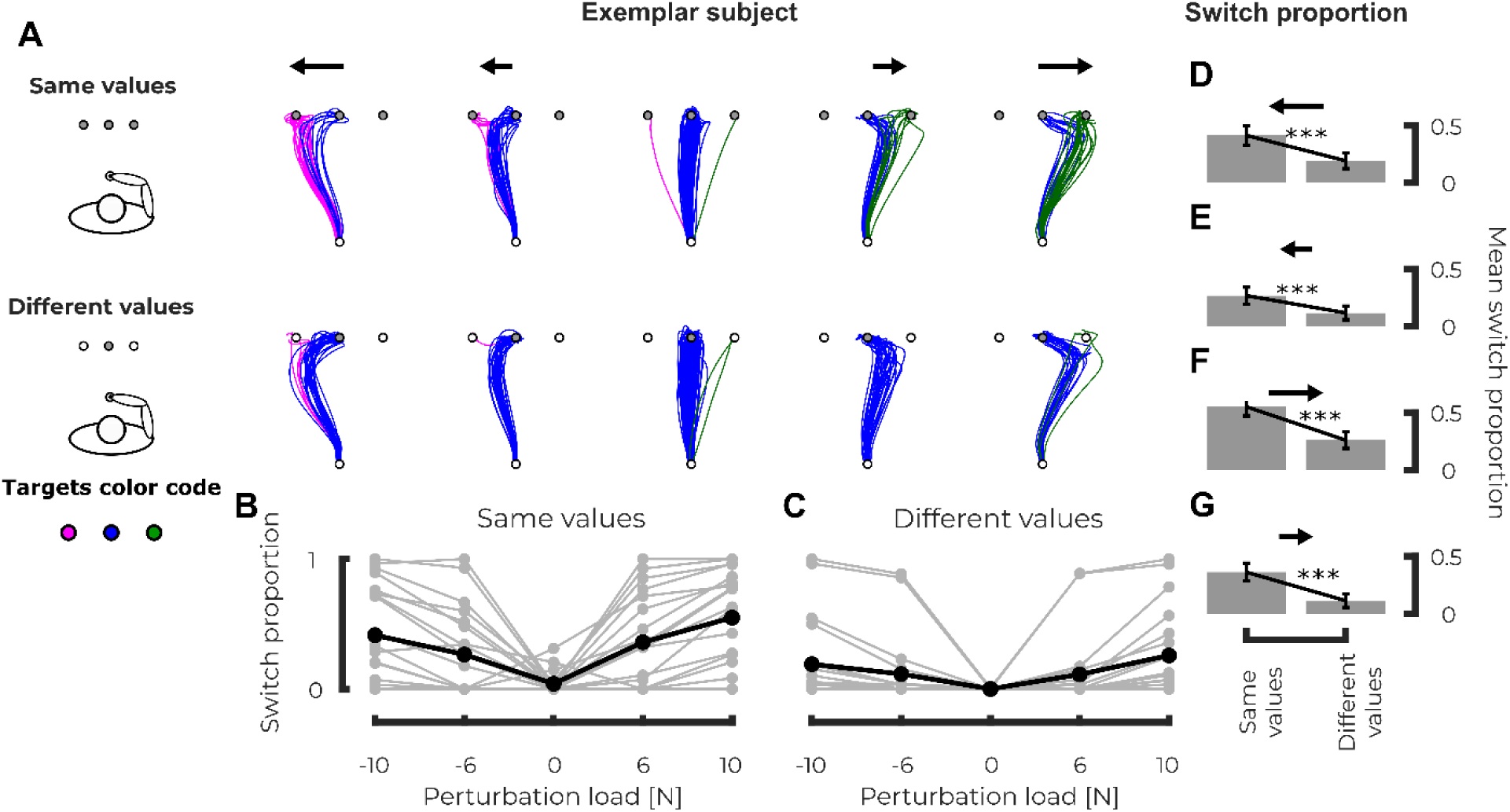
Experiment 2, Kinematics – **A** Representation of hand path of individual trials for a representative subject in the same condition on top (all the targets had the same reward) and in the different condition on bottom (the central target had a higher reward than the other two). The different columns represent the different force levels (from right to left, large leftward perturbation to large rightward perturbation). Magenta, blue and green paths respectively represent the paths that reached the left, central and right targets. **B** Group mean (black) and individual means (grey) of the switch proportion (i.e. fraction of trials that reached either the left or right targets) as a function of the applied load for the same condition. **C** Group mean (black) and individual means (grey) of the switch proportion (i.e. fraction of trials that reached either the left or right targets) as a function of the applied load for the different condition. **D-G** Comparison of the switch proportion for the same (left) and different (right) conditions for the trials with large leftward force, small leftward force, small rightward force and large rightward force respectively. p<0.05 (*), p<0.01(**), p<0.005(***)

These results showed that participants took the reward distribution of the options offered by the three targets into account while deciding which target they should reach. The next question that we will address is whether any parameters linked to the current state of the limb could modify the decision between the different motor outcomes.

Interestingly, we observed a link between the state of the limb at perturbation onset (kinematics and EMG activity) and the outcome of the motor decision. Figure 5A represents the mean EMG activity recorded in PM (top) and PD (bottom) in presence of mechanical perturbations (rightward, first column and leftward second column) across participants for the different targets (purple: left, blue: center and green: right) in the *same values* condition. No significant differences were observed in PM prior to force onset (−150ms to 0 ms, grey rectangle in Figure 5A) between the trials that reached the center target and the ones that reached the lateral targets (Figure 5B top) for both force directions (left: linear mixed model *β*_1_=-0.019, t=-1.76, p=0.0782, *r*^2^=0.62 and right: linear mixed model *β*_1_=0.0054, t=0.89, p=0.3758, *r*^2^=0.64). However, we observed an increase in the EMG activity of PD prior to perturbation onset for the trials that reached the center target compared to the ones that reached the lateral targets for both force directions (Figure 5B bottom, left: linear mixed model *β*_1_ = 0.022, t=3.78, p<0.005, *r*^2^=0.68 and right: *β*_1_=-0.051, t=-4.804, p<0.005, *r*^2^=0.60). This increase in EMG activity for trials that ended at the central target was correlated with larger forward velocities at force onset. We reported in Figure 5C the differences in forward velocities between the center and the lateral trials for both perturbation directions. In presence of a leftward perturbation (Figure 5C and D, right panels), we observed a larger forward velocity at force onset for trials that end up at the center target compared to those that reached the lateral target (linear mixed model: *β*_1_=-0.013, t=-3.347, p<0.005, *r*^2^=0.54). The same holds for trials with rightward perturbations (Figure 5C and D, left panels - linear mixed model: *β*_1_=0.040, t=9.476, p<0.005, *r*^2^=0.57). Similar observations were reported in the *different values* conditions. Indeed, we observed an increase in EMG activities in both muscles for the trials that ended up at the center target compared to those that reached the lateral target (PM linear mixed model: *β*_1_=-0.051, t=-4.81, p<0.005, *r*^2^=0.60 and PD linear mixed model: *β*_1_=0.022, t=3.78, p<0.005, *r*^2^= 0.68). Moreover, some tendencies were observed in the forward speed for trials with rightward (*β*_1_=-0.011, t=-2.019, p=0.0436, *r*^2^=0.52) and leftward (linear mixed model *β*_1_=0.012, t=1.95, p=0.0505, *r*^2^=0.57). These results collected in the *different values* conditions have to be analysed with caution because of the low number of trials that ended up at one of the two lateral targets (7.5% in the *different values* and 23.5% in the *same values* condition).

**Figure 5:**
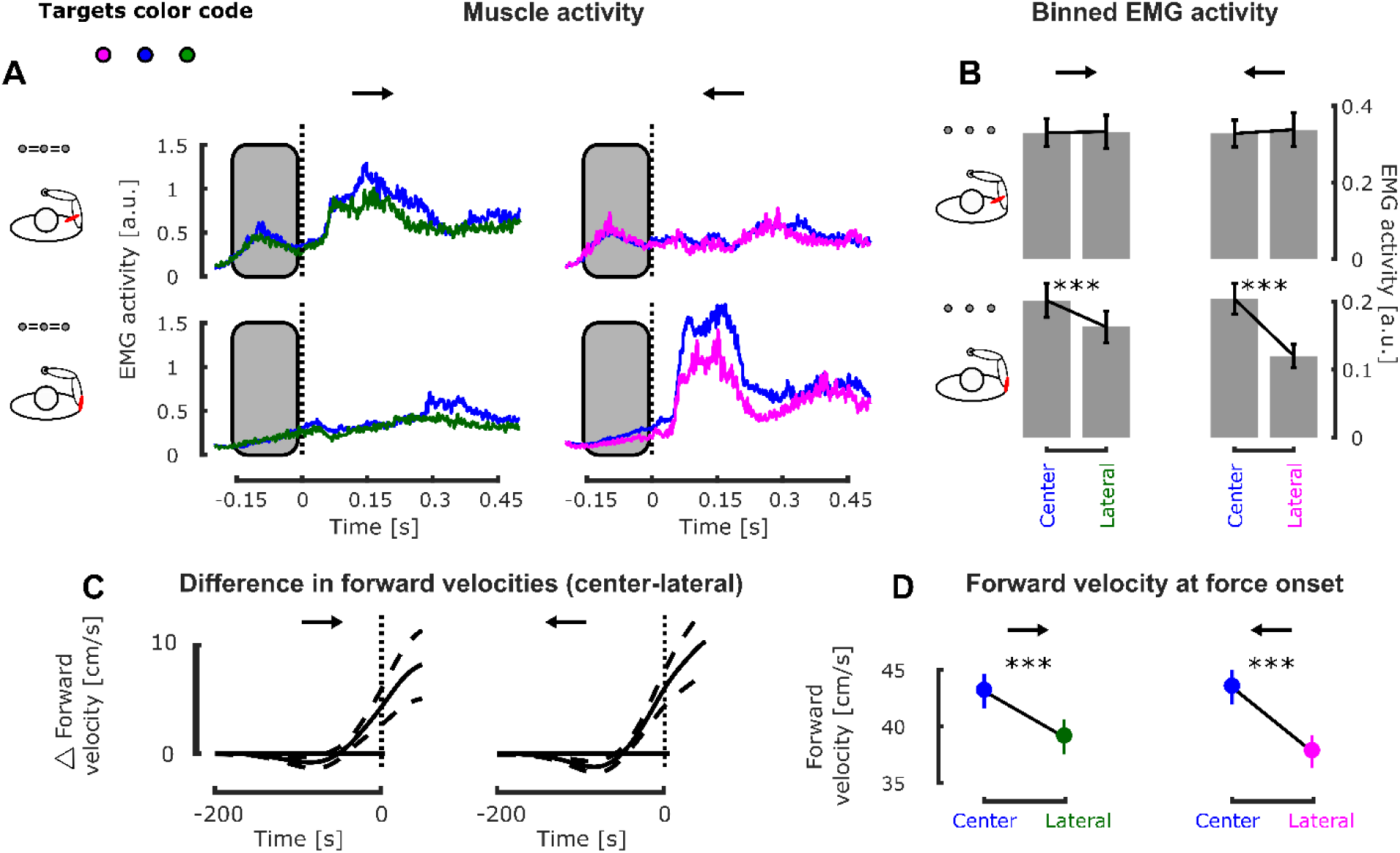
Experiment 2, EMG activity-**A** Mean EMG activity in Pectoralis Major (top) while responding to rightward (first column) and leftward perturbations (second column) and in Posterior Deltoid (bottom) while responding to rightward (first column) and leftward (second column) perturbations in the second experiment. The magenta, blue and green traces represent the mean EMG activity measured when participants reached the left, center or right target respectively. **B** Binned EMG activity before force onset in Pectoralis Major (top) and Posterior Deltoid (bottom) for the leftward and rightward perturbation loads, for the trials that reached the central (left bin) and lateral (right bin) targets. **C** Group mean and SEM of the differences in forward velocities across participants between the center and lateral trials for trials with rightward (left) and leftward (right) perturbations. **D** Comparison of the forward velocity at force onset for the trials that reached the central (blue) and lateral (green or purple) targets with a rightward or leftward perturbation load. p<0.05 (*), p<0.01(**), p<0.005(***)

We also tested whether the reward condition (ie. *same values* and *different values*) modified movement vigor by comparing the forward velocities and muscle activities at force onset between both reward conditions. We did not observe any difference in forward velocities between both reward conditions at perturbation onset as reported by mixed effect models (t=0.60, p=0.54, *r*^2^=0.03). Similarly, we did not observe any differences in EMG activities averaged during the 50ms preceding perturbation onset as revealed by mixed model analyses (PM: t=-0.3962, p=0.69, *r*^2^=0.28 and PD: t=0.07, p=0.93, *r*^2^=0.06). The same observation holds for reaction times that did not show any modulation with the reward condition (linear mixed model, p>0.05). This absence of correlation between the reward condition and movement vigor was interesting as it confirmed that we did not introduce any experimentally induced modulation of vigor in our paradigm. The differences in switching frequencies observed between the *same* and *different values* conditions are therefore attributable to the reward distribution and to vigor variability within both reward conditions.

This second experiment showed that humans take the rewards of the competing options into account to respond to perturbations and potentially change target goal during movement. More specifically, participants will tend to reduce their frequency of reaching toward targets that have a lower reward. We also showed that the state of the limb at perturbation onset modulated participants’ behaviour. Indeed, higher feedbacks gains at perturbation onset were correlated with a higher probability to potentially change target goal during movement, conditioned by the occurrence of mechanical load that would push the limb toward lateral target.

### Effect of the pre-activation of muscle on the motor decision

An outstanding question is when was the decision made to switch target. Did participants decide to change after the perturbation, or did they plan to change prior to movement? On the one hand, in this experiment as in previous reports (Nashed et al., 2014), changes in goal target depend on the occurrence and magnitude of the force so it is at least partially determined by sensory information collected during movement. On the other hand, the observation that the switch also depended on the baseline activity suggests that there could be an influence of the state of the limb from the beginning of the movement on the decision. We wanted to investigate this possibility in Experiment 3. This experiment was specifically designed to investigate whether the pre-activation of PD prior to movement onset could bias the frequency of target switches. Participants had to reach any of the three targets located at the same position as in Experiment 2. All targets had the same reward in this experiment. During movement, mechanical perturbation loads could push participant’s hand orthogonally to the main reaching decision. During half of the trials, a leftward background load was applied to participant’s hand throughout movement evoking a background activation to counter the background load (see Methods). We assessed the effect of pre-activation of PD by investigating the reach proportions to the lateral targets as a function of force intensity and background condition.

The application of a leftward background force induced an increase in both PM and PD baseline EMG activity (Figure 6C). We found a significant effect of the background load in both muscles (main effect of the linear mixed model on both muscles: *β*_1_=-0.11, t=-6.73, p<0.005, *r*^2^=0.91) as represented in Figure 6D. Moreover, we also observed an interaction effect between the background load and the muscle: baseline activity in PD increased more than PM activity (*β*_12_=0.20, t=19.067, p<0.005, *r*^2^=0.91).

**Figure 6:**
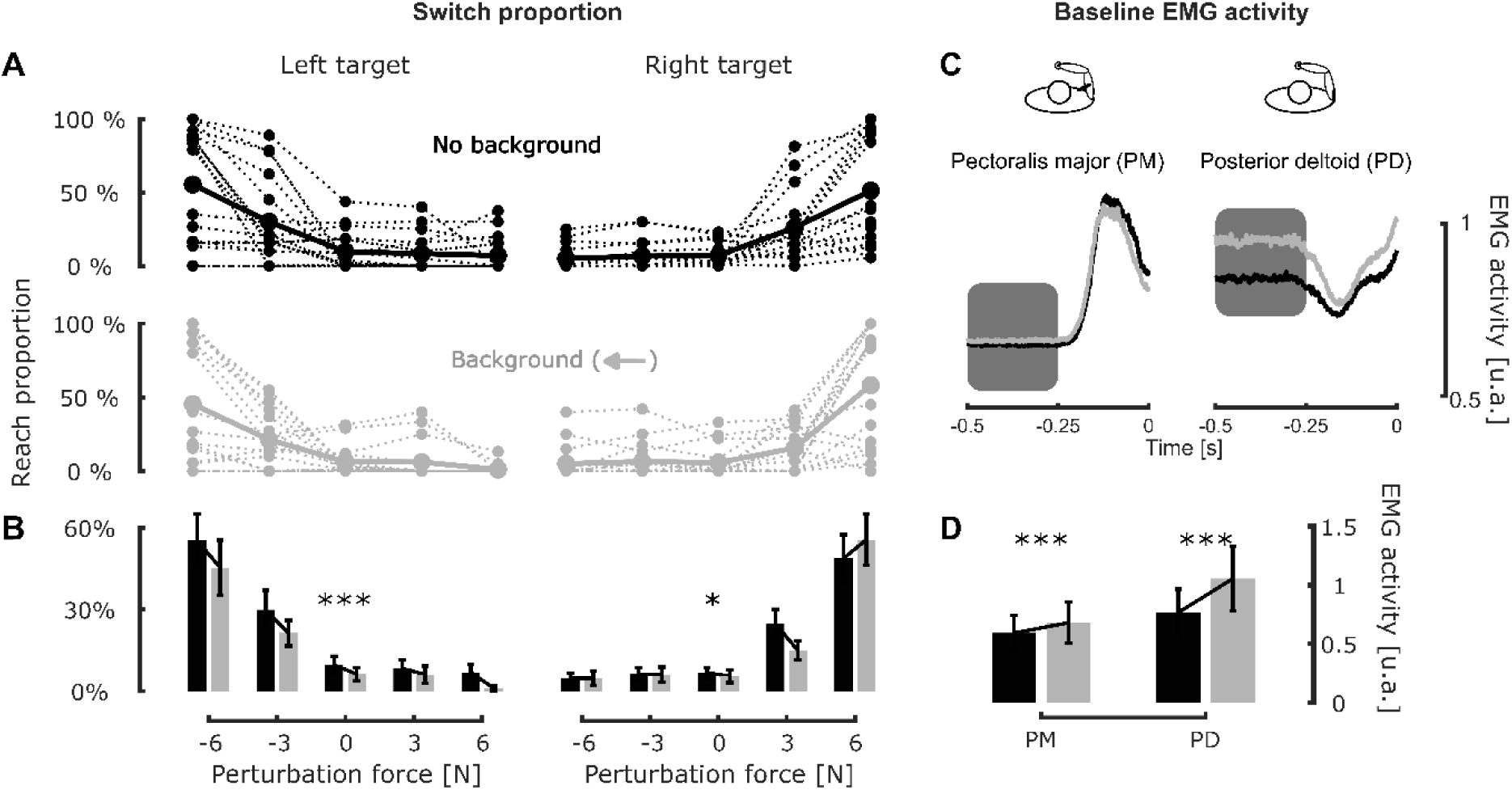
Experiment 3 – **A** Group mean (full line) and individual means (dashed lines) of the reach proportion to the left and right targets for the conditions without (top row, black) and with (bottom row, grey) the leftward background load as a function of perturbation load. **B** Comparison of the reach proportion of the left and right targets (left and right columns respectively) with (grey boxes) and without (black boxes) the leftward background force. **C** Group mean of the EMG activity in Pectoralis Major (left) and Posterior Deltoid (right) prior to movement onset for trials without (black) and with (grey) a background load. The time axis is aligned with the perturbation load onset. **D** Comparison of the binned EMG activity between 500 and 300ms (corresponding to the grey box in panel C) before force onset in Pectoralis Major and Posterior Deltoid for the conditions with and without background. p<0.05 (*), p<0.01(**), p<0.005(***)

We found that the leftward background load modified the reach proportion to the left target for all kind of online mechanical loads. Figure 6A represents the reach proportions to the left and right targets (respectively left and right column) as a function of the intensity of the perturbation load for the trial with (bottom) or without (top) background. In order to show the effect of the background load on the reach proportion to the lateral targets, we fitted a multilinear logistic regression (see Methods) that inferred the effect of perturbation and background load on the reached target. This multilinear logistic regression revealed a significant effect of the background and perturbations loads on both the left and right targets reaching proportion. Concerning the perturbation load, we observed an increase of the reach proportion to the left target with increasing leftward force (*β*_1_=- 0.9223, t=-22.16, p<0.005) and the mirror effect for the right target (*β*_1_=1.0204, t=23.86, p<0.005). The background load also had a significant effect on the reach proportion for these two targets. The reach proportion to the left target decreased when the background load was applied (*β*_1_=-0.4611, t=- 6.1759 and p<0.005, Figure 6B left panel). Intuitively an increase in force toward a target could bias the choice for that target but it was not the case. A slight decrease in reach proportion for the right target was also revealed by this regression (*β*_1_=-0.1544, t=-1.9972, p=0.0458, Figure 6B right panel). The intensity of this effect on the two lateral targets was compared using bootstrap resampling: this effect was larger for the left than for the right target. We generated 1000 bootstrap datasets from the original dataset used to fit the multilinear logistic regression and fitted the multilinear regression on each of these bootstrap datasets (generating that way estimates of *β*_1_ for each resampled dataset). We extracted bootstrap estimates of the main effect of background on the target reached for both lateral targets and computed the difference between the left and right estimates. The mean value of this difference was 0.280 and the 95% confidence interval obtained from bootstrap resampling was [0.072, 0.503] which therefore indicates a non-zero difference. This result suggests a directional bias in the effect of the background load on the switching strategies: the application of a leftward background load hindered switches to the left target more than those to the right one.

Post-hoc analyses performed on the individual reach proportion to lateral targets confirmed this asymmetry between left and right target (see Figure 6B). We observed a significant decrease of the individual reach proportion to the left target induced by the background load across participants and force levels (one tailed Wilcoxon signed-rank test : z=2.83, ranksum=999.5, p<0.005, d=0.21). No similar effect was observed for the right target (Wilcoxon signed-rank test: z=1.23, ranksum = 1015, p=0.2154, d=0.03).

An interesting question is whether this background force also modulated forward velocity. We address this question by using a linear mixed model to compare forward speed at force onset in the conditions with and without background load. No modulation of movement speed between these conditions was observed (linear mixed model *β*_1_=-0.009±0.007, t=-1.2528, p=0.210, *r*^2^=0.44). This result is important as it discards the eventuality that the modulation of flexibility to switch to a new target goal was induced by movement velocity. Similarly, the reaction time was not modulated by the presence of the background force (linear mixed model, p>0.05).

The results of this last experiment showed that the tendency to switch observed in Experiment 2 depended on the biomechanical state of the limb. Importantly, the application of a background load in a direction reduces the tendency to switch in this direction in a larger amount than the tendency to switch in the opposite direction.

## Discussion

We conducted a series of experiments to investigate the impact of reward on feedback control strategies and rapid motor decisions by probing the impact of explicit target reward. In Experiment 1, we demonstrated that target reward does not only increase movement vigor as reported in previous studies (Summerside et al., 2018; Yoon et al., 2018), but it also increases feedback responses and muscle activity. We observed that perturbation-related lateral hand deviations were smaller when participants reached toward a target associated with higher reward. Moreover, we also observed an increase in the baseline EMG activity as well as an increase in the EMG responses to perturbation loads with increasing reward. Altogether, these results suggest that the feedback gains used to perform movements scale with the value of the reward. In the second experiment, we reported that the reward distribution across the competing options has an influence on rapid motor decisions: participants were less prone to switch to a nearby target if it was associated with lower reward. In addition, an increase in feedback gains was detrimental to the ability to switch target during movement as we observed in Experiments 2 and 3 that participants were also less likely to switch target during movement when the muscle activity was higher. The modulation in muscle activity introduced experimentally in Experiment 3 induced a directional bias in the ability to switch target online, demonstrating a causal influence of muscle activity.

The increase in movement vigor and feedback gains associated with reward that we observed in Experiment 1 was coherent with the selection of a robust control strategy (Bian et al., 2020; Crevecoeur et al., 2019). A robust controller consists in an alternative to stochastic optimal control (Todorov & Jordan, 2002) that has the property to consider unmodelled disturbances (Basar & Bernhard, 1991) which results in better responses to mechanical perturbations during movements. Reward is known to invigorate movements as revealed in saccadic eye movements where faster movements were observed toward higher monetary rewards (Manohar et al., 2015, 2017) or toward targets associated with higher implicit rewards (Xu-Wilson et al., 2009). Similar observations were made for upper limb reaching movements that exhibited higher peak velocities toward more rewarding targets (Esteves et al., 2016; Sackaloo et al., 2014; Summerside et al., 2018; Yoon et al., 2018). This was taken as evidence for reward-dependent selection of movement time (Haith et al., 2012; Shadmehr et al., 2010). It has recently been demonstrated that this increase in movement vigor was associated with higher muscle activity in presence of reward (Codol et al., 2020) which could be interpreted as a mechanism used to increase internal feedback gains in order to improve reward-related endpoint accuracy (Manohar et al., 2019). Here we postulate that another mechanism is also at play: a higher reward produced a more robust strategy that revealed participants’ will to render their movements less sensitive to perturbations, thereby reducing the risk to miss the goal. In this framework the reduction in movement time results from the robustness of the control that impacts movement velocity through larger goal directed control gains.

In this framework, the modulation of the robustness of control has a clear limitation that we were able to establish empirically: a robust control strategy is meant to reject disturbances indistinguishably, thus in principle it is clear that this strategy is not compatible with a flexible change in movement goal online, which requires a reduction in feedback response to let the perturbation redirect one’s hand toward the new goal.

Besides the property of the robust model to predict larger feedback gains, we measured here as in previous work that this strategy was associated with an increase in baseline co-activation (Crevecoeur et al., 2019) which potentially influences the gains of short- and long-latency responses to mechanical perturbations (Bedingham & Tatton, 1984; Marsden et al., 1976; Matthews, 1986; Pruszynski et al., 2009; Stein et al., 1995; Verrier, 1985). Considering this, the competition between robust control and flexible online decisions in the human motor system may depend in part of the fact that the robust controller recruits the peripheral motor apparatus (i.e. muscle state and reflex gains) to increase the overall feedback gains, thereby creating a competition between peripheral mechanisms engaged in control and more central decisional processes.

Moreover, our results demonstrate that the model based on distributed consensus of decision-making (Cisek, 2012) also applies to online motor decisions. This framework posits that decisions occur through an competition between the different options by integrating the motor costs incurred to each action (Cos et al., 2011; Morel et al., 2017; Shadmehr et al., 2016) and their respective outcome (Trommershäuser et al., 2003, 2008). We documented a concomitant influence of both the reward distribution across competing options and load magnitudes which highlights that these two factors are taken into consideration during online motor decision. In addition, we add to these factors that the state of the peripheral motor system, influenced by the selected control strategy and feedback gains, had an effect on online decision-making. Our findings are in line with previous work reporting an impact of the cost of each action (Kurtzer et al., 2020; Michalski et al., 2020; Nashed et al., 2014) and their associated outcome (Cos et al., 2021; Marti-Marca et al., 2020). These observations support that online motor decisions must result from distributed consensus between control strategies, feedback responses and rewards. Importantly, the present study investigated participants’ decision to switch to alternative targets during movement, all of which leading to successful movements, there were no good or bad choices as it is the case in a *go-before-you-know* paradigm (Chapman et al., 2010; Enachescu et al., 2021; Gallivan et al., 2016, 2017).

To conclude, our study highlights that multiple mechanisms underlie reward-dependent planning and control of movement. One the one hand, we suggest that there is a robust control strategy that involves peripheral circuits by means of increases in baseline activity and gain scaling of the feedback responses. This strategy associated with robust control is likely selected to reject perturbations and reduces the risk of missing the reward suggesting that there could be a cost incurred to reward. On the other hand, there exists a more flexible control strategy able to switch target during movement. It is conceivable that this second strategy, which requires some inhibition of muscle activity and response, is mediated by higher level inhibitory circuits and response modulation (Scott, 2016; Shadmehr & Krakauer, 2008). Both strategies integrate explicit target rewards and depend on the state of peripheral control loops.

An interesting open question is whether and how much individuals can modulate their strategy or whether the differences in strategy reflect individual traits. Indeed, individual differences have been shown in movement vigor (Reppert et al., 2018), and their possible effect on the modulation of feedback control is an exciting open question. Such ability is potentially central to understand planning and control in complex environments.

## Notes

### Competing Interest Statement

The authors have declared no competing interest.

